# Differential expression of sex regulatory genes in gonads of *Astyanax mexicanus* surface fish and cavefish

**DOI:** 10.1101/2024.12.22.629934

**Authors:** Kaitlyn A. Webster, Bethany Ponte, Hans Vasquez-Gross, Juli Petereit, John Hutchinson, Misty R. Riddle

## Abstract

**Background:** *Astyanax mexicanus* is a single species of fish that consists of river-dwelling (surface) and cave-dwelling morphotypes. Little is known about how sexual determination, differentiation or reproduction have evolved in the surface morphs or cavefish, though divergence in reproductive strategy is expected as the latter have adapted to the novel cave environment. Evolution of the gonad transcriptome may underlie the differences in gamete morphology, fertility, and fecundity previously reported between morphotypes.

**Results:** We compared the ovary and testis transcriptome of surface fish and cavefish at juvenile and adult stages. We found that samples clustered by developmental stage, sex, and morphotype identity. Several key genes that are typically associated with the female gonad in other vertebrates showed a reversal in sexual dimorphism or were not differentially expressed between sexes in *A. mexicanus*. In contrast, while gene expression typically associated with male gonads was largely conserved and consistent with vertebrate testicular expression profiles. Transcriptional and physiological differences between surface fish and cavefish morphotypes were observed in gonads from both sexes. Cavefish ovaries exhibited unique upregulation of neuron development and differentiation genes, and extensive innervation of the ovarian epithelium, while cavefish testes showed increased expression of angiogenesis regulating genes, and greater vasculature density compared to surface fish testes.

**Conclusions:** These results reveal significant gene expression differences between *A. mexicanus* surface fish and cavefish morphotypes that may have functional consequences in gonad morphogenesis and fertility. Our findings provide a foundation for investigating the evolution of sex regulatory pathways and reproductive strategies in animals adapting to new and challenging environments in which nutrient availability, temperature, and mate selection are suboptimal.

## Background

Studying gonadal sex differentiation in fishes can help identify novel molecular pathways that mediate sex determination and development. In contrast to mammals, which universally utilize chromosomal XY sex determination directed by the *SOX3*-derived *SRY* gene, fishes display a diversity of sex determining mechanisms (SDM) and sex regulatory genes. Across teleosts, SDM are known to be plastic, rapidly evolving, and inclusive of all known modes of regulation: chromosomal, polygenic, hormonal, social, and environmental (Devlin and Nagahama 2002; Schartl 2004). Different SDM and master sex regulatory genes can evolve even within the same genus, as reported in closely related species of Medaka rice fish (Matsuda 2002, Myosho 2012, Takehana 2014). Like other non-mammalian vertebrates, teleost fish sex determination can be influenced and overridden by social dynamics and environmental conditions, even when sex chromosomes or sex regulatory genes are employed. Notably, environmental factors such as photoperiod, temperature, salinity, pH and hypoxia have been shown to skew sex ratios in teleosts (Brown 2014, Baroiller 2009, Abucay 1999, Shang 2006). Despite the diversity of SDM, most vertebrate sex regulatory genes have orthologues in teleosts, which have been well studied in zebrafish, Medaka, and tilapia. For example, the mammalian ovary-promoting genes *Foxl2*, *Wnt4*, and *Cyp19* (aromatase) show conserved female-specific function in teleosts, as do the testis-promoting genes *dmrt1*, *amh*, and *sox9* (Yang 2017, Kossack 2017, Lau 2016, Guigen 2010, Webster 2017, Yan 2019, Nakamura 2012). How sex differentiation and gonad development evolve as fishes adapt to new environments is not well understood.

The Mexican tetra, *Astyanax mexicanus,* is a species of fish that consists of river-adapted and cave-adapted morphotypes. There are as many as 32 geographically separate cave-dwelling “cavefish” populations that evolved from river-dwelling “surface fish” in the underground limestone caves of the El Abra Region of Tamaulipas and San Luis Potosí, Mexico (Figure 1D) (Espinasa et al., 2020). Based on whole genome sequencing of field-collected individuals, the *A. mexicanus* phylogeny defines two lineages; cavefish from the Tinaja and Pachón caves form a monophyletic clade separate from the Molino cavefish and Río Choy surface fish (Herman et al., 2018). This evolutionary history indicates that cavefish traits have evolved through repeated evolution providing researchers with natural replicates to investigate the genetic and developmental basis of cavefish traits.

**Figure 1.**
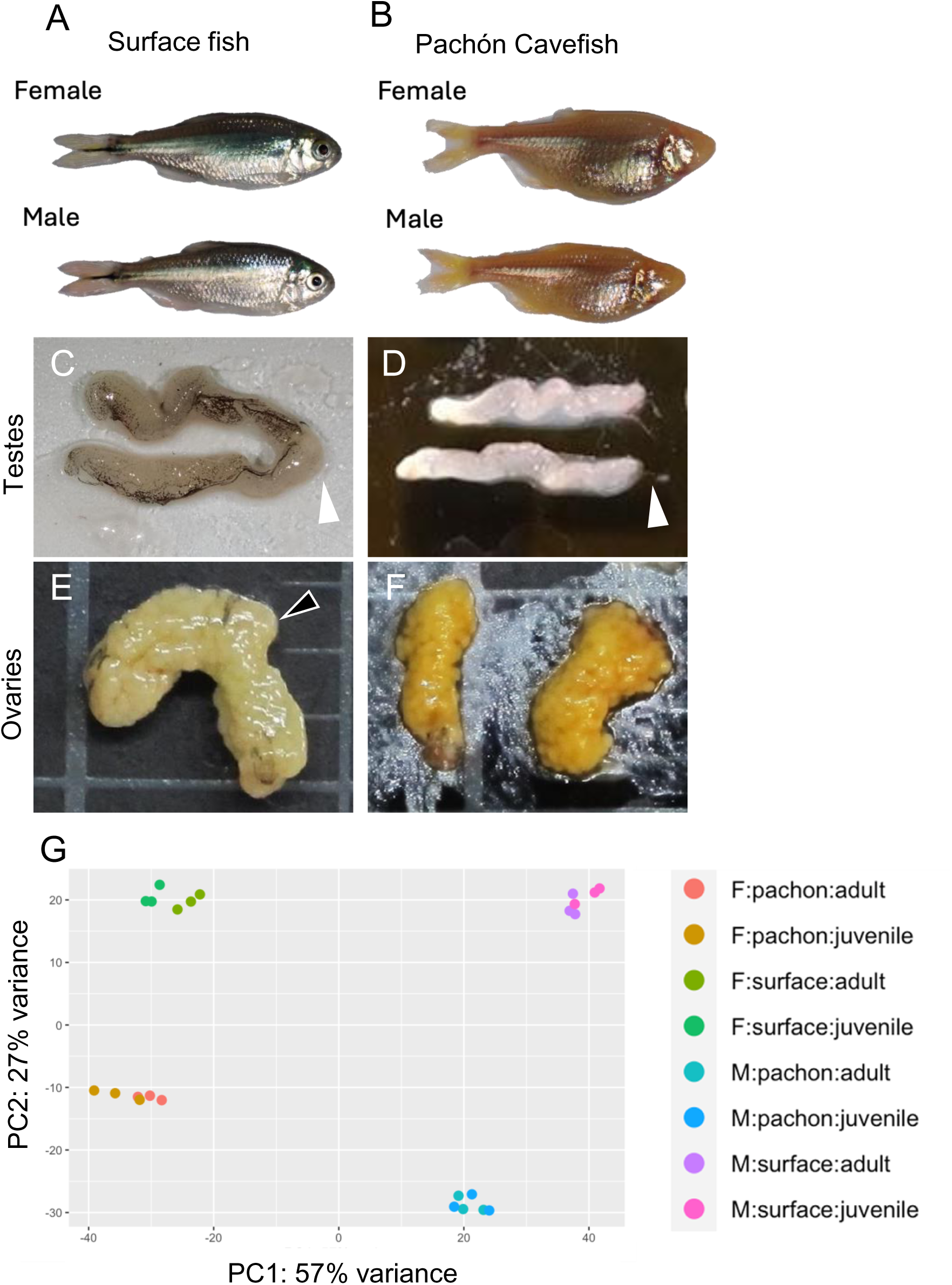
Transcriptome analysis of *Astyanax mexicanus* gonads. Appearance of *A. mexicanus* surface fish (**A**) and Pachón cavefish (**B**) males and females. Appearance of sexually mature testis and ovaries of surface fish (**C, E**) and Pachón cavefish (**D, F**). Testes (**C, D**) consist of two lobes of tubules containing male germ cells and sperm and terminating at the efferent urogenital duct (white arrowheads). Ovaries (**E-F**) consist of two lobes containing clusters of female germ cells and eggs surrounded by the germinal epithelium and terminating at the oviduct (black arrowhead). (**G**) Principal component analysis (PCA) plot of normalized, variance stabilized RNA sequencing expression profiles of *A. mexicanus* gonads. Each point represents an individual sample, colored by sample type. The first principal component (PC1) explains 57% of the total variance and separates samples by sex. The second principal component (PC2) explains 27% of the total variance and separates samples by morphotype (surface fish vs Pachón cavefish). PCA plots based on differentially expressed genes (DEG) only (adjusted p-value less than 0.05).

Little is known about sex and gonad development in *A. mexicanus* despite its growing importance as a model system in understanding the genetic basis of evolution (Perera 2023). For example, only a few studies have compared the expression or function of sex-associated genes between surface fish and cavefish morphs. In Pachón cavefish, sex-associated genes such as *foxl2a*, *wnt4b*, *cyp19a1a*, *dmrt1*, and *amh* were reported to be expressed in the gonads but surprisingly *foxl2a*, *wnt4b*, and *cyp19a1a* were not found to be significantly sexually dimorphic (Imarazene 2021a). Pachón cavefish male sex is determined by supernumerary “B-sex” chromosomes that exhibit non-mendelian inheritance (Imarazene et al., 2021b). The gene conferring male sex on the Pachón cavefish B chromosome is *growth differentiation factor 6b* (*gdf6b*) which if knocked out results in male-to-female sex reversals (Imarazene et al., 2021b). The mechanisms of sex determination in surface fish and other cavefish populations have yet to be described. The atypical expression of conserved vertebrate sex-specific genes and emergence of a novel sex determination mechanism in Pachón cavefish highlights the promise of this model for the study of evolution of sex regulatory pathways and adaptive reproductive strategies. In addition, since traits like body condition and relative organ size are sexually dimorphic and are controlled by loci that have sex-dependent effects (Riddle et al., 2021), revealing how sex determination and gonad development differs between *A. mexicanus* morphs is crucial to understanding cavefish evolution.

Here, we describe the first transcriptomic analysis of ovaries and testis in *A. mexicanus,* comparing surface fish and Pachón cavefish at juvenile and adult stages. We found that samples clustered by developmental stage, sex, and morphotype identity. We found that in surface fish and Pachón cavefish, typical female-associated genes are not differentially expressed between males and females. However, typical male-associated gene expression is conserved. We observed an upregulation of genes involved in neuron development in Pachón cavefish ovaries compared to surface fish ovaries and found that the germinal epithelium surrounding germ cells is extensively innervated in both morphotypes. Comparing the testis, we observed increased expression of angiogenesis genes in Pachón cavefish compared to surface fish and evidence of more vasculature within the Pachón cavefish testis. Our results provide insight into the molecular, functional, and structural differences between surface fish and cavefish gonads which is essential for future studies examining the evolution of sex differentiation, sexual dimorphism, and reproduction in this model system.

## Methods

### Animal husbandry

Río Choy surface fish and Pachón cavefish were derived from parents bred for greater than five generations in the laboratory of Dr. Clifford Tabin at Harvard Medical School. All animals used in this study were maintained, housed, and fed under the same conditions on a recirculating water system (Temperature: 23-24 °C, pH: 7–7.5, Conductivity: 1200-1400μS) in a facility with a 10:14-h light/dark cycle. Adult fish were fed with a combination of New Life Spectrum TheraA+ small and/or medium sinking pellets (dependent on fish size) and *Artemia* and housed at a density equal to two adult fish per liter of water. Hatched larval fish were fed rotifers up to 14 days post fertilization (dpf) in 1-liter cups of fish-ready water, then transferred to the recirculating system where they were fed *Artemia* through 60 dpf. At 1.5 years post fertilization, fish were euthanized in MS-222 prior to gonad dissection. We found that adult Pachón cavefish males and females were on average larger than surface fish counterparts with larger gonads, but did not observe significant differences in size between morphotypes (Pachón cavefish versus surface fish average fish weight: male = 1.34g vs 0.95g, p = 0.09, female = 1.9g vs 1.2g, p = 0.17, t-test, Pachón cavefish versus surface fish average gonad weight: testes = 0.034g vs 0.024g, p = 0.40, ovaries = 0.165g vs .097g, p = 0.46, Additional file 1).

### Sample collection, RNA extraction and cDNA synthesis

Juvenile (i.e., not containing ovulatory eggs or caudal duct spermatozoa) to adult fish aged between 1-1.5 years were euthanized in 400ppm MS-222. The gonads were dissected into cold 1 X PBS, and thoroughly cleared of associated fat or viscera using a dissecting microscope.

Cleaned gonads were then collected into 0.5mL (adult gonads) or 0.25mL (juvenile gonads) Trizol and homogenized using a hand-held pellet pestle, then stored at −80°C. Total RNA was extracted using Zymo Research Direct-zol RNA MicroPrep with DNAse treatment according to the manufacturers protocol. LunaScript RT Supermix kit with 1μg of RNA was used to synthesize cDNA. Diluted cDNA samples (50ng/μL) were used for sequencing.

### RNA sequencing and gene ontology analysis

Illumina sequencing was performed by the Nevada Genomics Center using NextSeq 2000 P3 100 cycle sequencing kit. Samples were sequenced with 50bp paired-end reads with a mean of 52.2 million reads (minimum depth of 29.6 to 58.8 million reads per sample). FastQC v0.11.9 was ran on each sequenced sample both pre and post read trimming to assess improvement after trimming. Trimmomatic v0.36 with parameters ILLUMINACLIP:2:30:10 SLIDINGWINDOW:4:15 MINLEN:36 was used in order to remove Illumina sequencing adapters and poor base quality regions.

Reads were aligned to Ensembl build 2 of the *A. mexicanus* genome, augmented with transcript information from Ensembl release 2.0.97 using STAR v2.7.5c (Dobin et al., 2013). STAR output alignment BAM files were imported into featureCounts v2.0 from the subread package in order to quantify reads per sample for genes. Quantified counts generated via featureCounts were imported into the statistical software package R (R-core-team, 2019) for downstream analysis. A low count was defined by less than 10 counts per gene. If a gene demonstrated low counts across all samples and experimental groups, it was removed from further downstream analysis. Differential gene analysis was performed with the DESeq2 package (Love 2014) and statistically significant differences were defined by a Benjamini-Hochberg FDR adjusted *p*-value <0.05. Variance stabilizing transformation (VST) was used to visualize count data in a Principal Component Analysis plot.

To gain more insight into the differentially expressed genes, gene names were annotated using the biomaRt package (Durinck 2005), and we determined enriched Biological Processes Gene Ontologies (BP GO) using clusterProfiler (Yu 2012). Gene ontology maps were generated from enriched Biological Processes GO terms for differentially expressed genes (padj < 0.05) using clusterProfiler cnetplot function (Yu 2012). Annotation hub AH107459 was used for *Astyanax mexicanus* GO term annotation.

### Ovary immunostaining

Ovaries from adult Pachón cavefish and surface fish were dissected, fixed in 4% PFA, and dehydrated in ethanol. After rehydration, the tissue was incubated with α-acetylated tubulin (1:500, Sigma-Alrich, T6793) in blocking solution (BSA, sodium azide, goat serum, DMSO, PBS, and Triton X-100) for 48 hours at room temperature, rinsed and incubated with a secondary antibody (goat anti-mouse 488 IgG H&L, ABCam, ab150117) for another 48 hours at room temperature. The tissue was imaged on a Leica THUNDER model organism microscope.

### Testes Vascular Staining and Quantification

Testes were dissected from adult Pachón cavefish and surface fish and collected into 1 x PBS at room temperature, briefly. Whole testes were then immediately incubated for 5 minutes in the dark in a solution of o-Dianisidine (3,3′-dimethoxybenzidine, MilliporeSigma D9143-5G) formulated as described in O’Brien 2009. Stained testes were dehydrated, cleared in xylene, and embedded in paraffin. Sections of 10 uM were then counterstained with modified Harris hematoxylin solution (Sigma-Aldrich HHS32). Tissue area was calculated using ImageJ. Each bundle of vascular staining was traced, and the area was recorded. The vascular area was summed and divided by the total tissue area in the image to calculate percentage of testicular vascularization.

## Results

We compared the ovary and testes transcriptome of *A. mexicanus* surface fish and Pachón cavefish that were 14-18 months post fertilization (Figure 1 A-E). Interestingly, regardless of chronological age, two distinct morphological stages of gonads were observed for each sex, referred to in this study as juvenile (not containing mature gametes) or adult (containing mature gametes). Gonads categorized as juvenile appeared small and translucent upon dissection, and by histology were confirmed to lack ovulatory oocytes (females) or spermatozoa (males). Adult gonads were large and opaque, and stored mature gametes in the urogenital ducts. Overall, the gonad morphology was similar comparing surface fish and Pachón cavefish, with the most striking difference observed in coloration. Surface testes were sheathed in scattered black melanocytes (Figure 1B), whereas Pachón testes lacked any melanin pigmentation consistent with albinism (Protas 2006) (Figure 1C). Conversely, Pachón cavefish ovaries were intensely yellow-orange compared to surface fish ovaries, consistent with increased carotenoid accumulation in the eggs (Riddle 2020) (Figure 1E).

We first visualized transcriptome differences between sample types (n=3 per developmental stage, sex, and morphotype) using principal component analysis and found that samples clustered according to sex and morphotype identity (Figure 1J). We found that PC1 (57% variance) separates testes from ovaries, while PC2 (27% variance) separates surface fish from Pachón cavefish. Juvenile and adult gonads formed overlapping clusters except for surface fish ovaries that showed no overlap between developmental stages. These findings indicate differential regulation of gene expression between *A. mexicanus* morphotypes within the gonads of males and females.

We examined the gene expression profiles that were specific to each sex to investigate if the function of known regulators of sex differentiation in other species are likely conserved in *A. mexicanus.* We found 17,955 differentially expressed genes (DEG) between the testes and ovaries of *A. mexicanus* (Figure 2A). We observed expected sexually dimorphic expression of some highly conserved and commonly studied sex regulatory genes, but also found that several genes which are customarily restricted to the ovary or upregulated in the ovary, as best characterized in the zebrafish model, are not sexually dimorphic (Figure 2B). For example, the germ-plasm aggregator bucky ball (*buc*) is more highly expressed in the ovary but is also unexpectedly abundant in the testes which do not express *buc* in other teleosts (Figure 2J). Other ovary-associated genes such as the estrogen synthesis enzyme aromatase (*cyp19a1a*), the gonadal somatic cell transcription factor Forkhead box protein L2 (*foxl2a*), and the secreted growth factor wingless-type MMTV integration site family, member 4 (*wnt4a*) were either not sexually dimorphic in the gonads (*cyp19a1a*, Figure 2C), or exhibit a sex-reversed expression profile, i.e. were unexpectedly more highly expressed in the testes (*foxl2a* and *wnt4a*, Figure 2D-E). These results suggest possible neofunctionalization of these sex regulators in *A. mexicanus*. In contrast to female-promoting genes, male-promoting genes displayed expected patterns of expression comparing ovaries and testes. For example, the TGF-β superfamily member anti-Müllerian hormone (*amh* Figure 2F), the androgen synthesis enzyme Cytochrome P450, family 11 (*cyp11c1*, Figure 2G), the doublesex and mab-3 related transcription factor 1 (*dmrt1*, Figure 2H), and the teleost-specific TGF-β superfamily member gonadal soma derived factor (*gsdf*, Figure 2I) were upregulated in the testes of both Pachón cavefish and surface fish.

**Figure 2.**
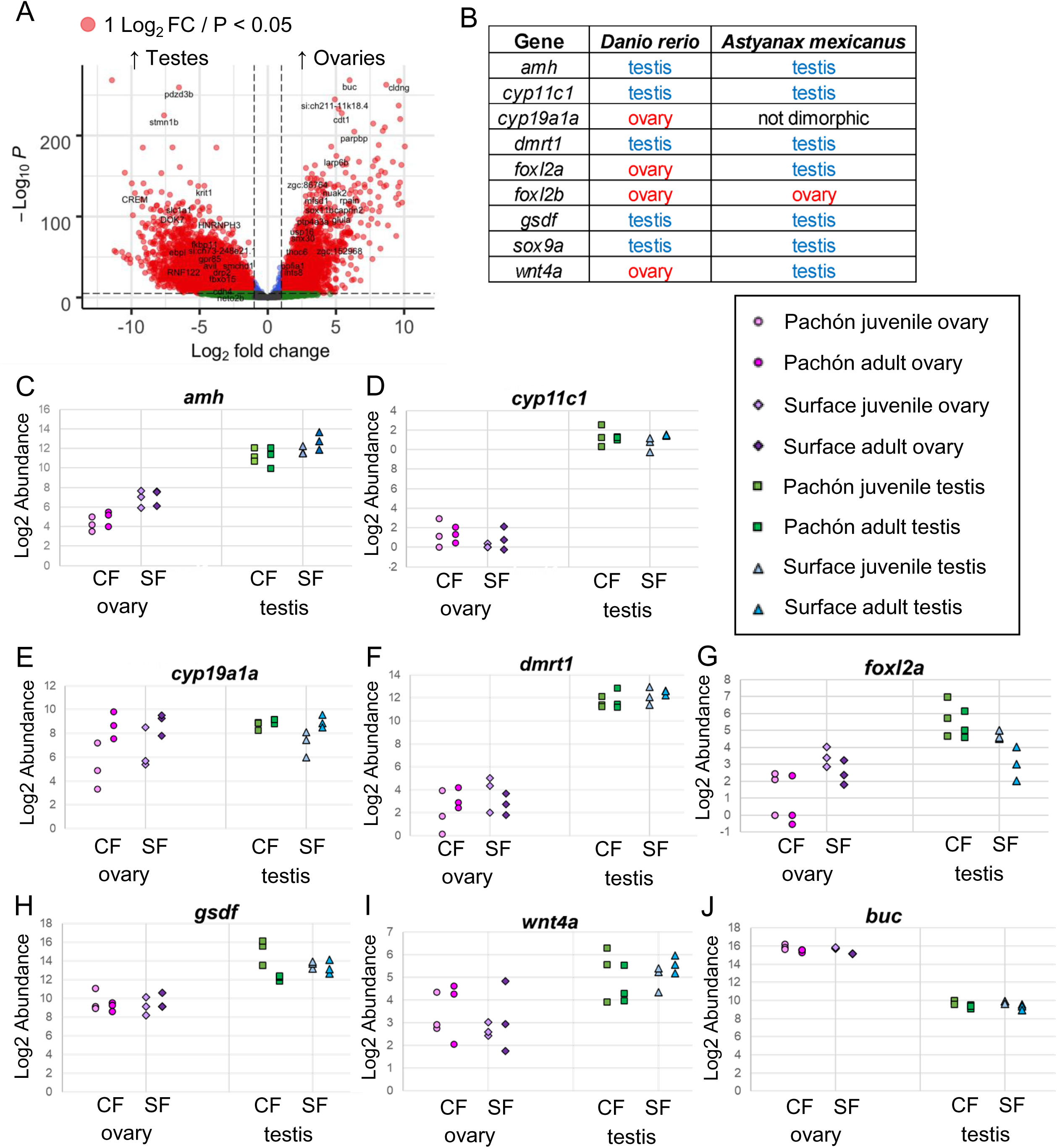
Sex-specific gene expression profiles in *Astyanax mexicanus* surface fish and Pachón cavefish gonads. (**A**) Volcano plot showing genes that are differentially expressed between *A. mexicanus* testes and ovaries (includes data from juvenile, adult, surface fish, and Pachón cavefish gonads). The y-axis shows −log10 adjusted p-value and the x-axis shows log2 fold change for all genes after low-count filtering (23,809 transcripts). Genes with adjusted p-value < 0.05 and absolute log2 fold change > 1 are highlighted in red. Negative fold change indicates testis-enriched expression, while positive fold change indicates ovary-enriched expression. (**B**) Summary of genes with sex-specific expression profiles in the gonads of zebrafish (*Danio rerio*), compared to the expression profiles we observed in *A. mexicanus.* “Ovary” indicates a gene with upregulation in female gonads, whereas “testis” indicates a gene with upregulation in male gonads. (**C-J**) Plots showing *A. mexicanus* gonad log2 transcript abundance of genes with sex-specific expression profiles and functions in the gonads of other vertebrates. Each point represents the transcript abundance in an individual sample, colored and organized on the x-axis by the sample type. From left to right: cavefish (CF) juvenile ovary, CF adult ovary, surface fish (SF) juvenile ovary, SF adult ovary.

We next compared the ovary transcriptome between surface fish and cavefish morphotypes (Figure 3). Comparing juveniles, we observed 7,312 DEG between morphotypes (Figure 3A). Interestingly, the gene that is most statistically significantly upregulated comparing juvenile Pachón cavefish versus surface fish ovaries was *spock2* (Testican-2), which is a calcium-binding proteoglycan that mediates oocyte maturation in mice. Comparing adults, we found 3,698 DEG (Figure 3B). We found that the top 3 gene ontology (GO) term categories for genes upregulated in Pachón juvenile ovaries are protein phosphorylation, neuron development, and plasma membrane bounded cell projection organization (Figure 3C). Similarly, in Pachón adult ovaries the top categories are nervous system development, neuron differentiation, and generation of neurons (Figure 3D). *Semaphorin 3aa* (*sema3aa*) is one of the most upregulated genes in Pachón ovaries of both stages and is implicated in the migration of specialized neurons that secrete gonadotropin-releasing hormone (GnRH) – a critical mediator of puberty and subsequent reproductive cycling (Giacobini 2015). Overall, these findings indicate an upregulation of neuron-associated genes in the ovaries of Pachón cavefish compared to surface fish.

**Figure 3.**
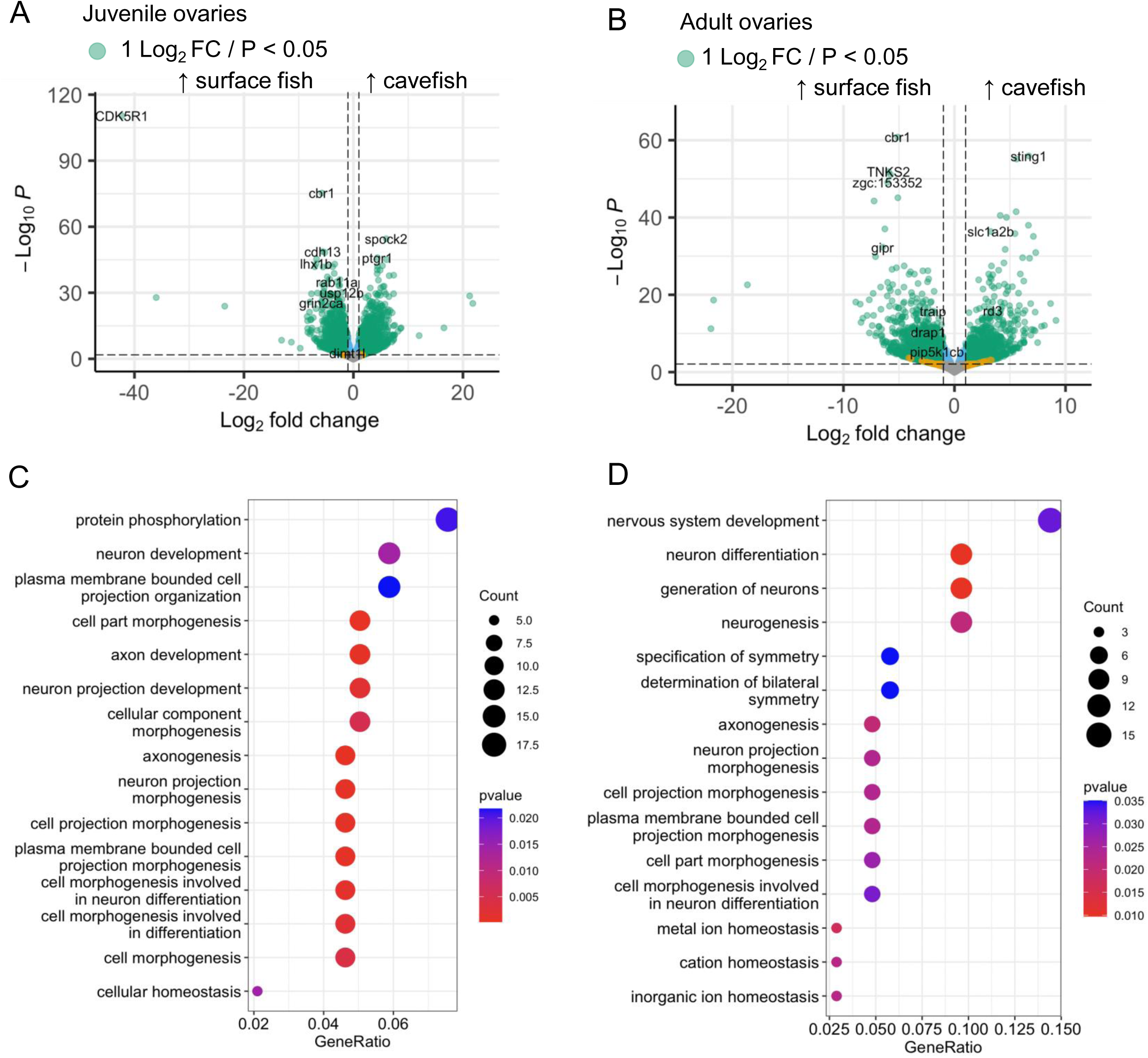
Ovary transcriptome differences between surface fish and Pachón cavefish suggest enhanced neurogenesis in cavefish. (**A, B**) Volcano plots showing genes that are differentially expressed between the ovaries of *A. mexicanus* surface fish and Pachón cavefish at juvenile (**A**) and adult (**B**) stages. The y-axis shows −log10 adjusted p-value and the x-axis shows log2 fold change for all genes after low-count filtering. Genes with adjusted p-value < 0.05 and absolute log2 fold change > 1 are highlighted in green. Negative fold change indicates greater expression in surface fish, while positive fold change indicates greater expression in cavefish. (**C**, **D**) Dot plots displaying top 15 enriched biological process gene ontology (GO) terms identified from gene set enrichment analysis of genes upregulated in Pachón cavefish compared to surface fish at juvenile (**C**) and adult (**D**) stages. Each dot represents a biological process GO category, with the size of the dot corresponding to the number of genes in the category, and the color scale indicating the adjusted p-value. The x-axis represents the gene ratio (proportion of input genes involved in the pathway), while the y-axis lists the enriched pathways.

In line with a role for neurons in the regulation of ovary function, we observed substantial innervation throughout the sheath that surrounds the ovary, the geminal epithelium, in both morphotypes (Figure 3F-H). A large neuron trunk that branches to reach throughout the ovary germinal epithelium was visible on medial side of the ovary while the lateral side of the ovary shows little to no innervation. A similar innervation pattern is observed in zebrafish and is thought to be important to regulate oocyte quiescence through norepinephrine signaling (Kim 2021). Our findings suggest that altered neuronal signaling may be important for the evolution of cavefish ovaries.

We next compared the testes transcriptome between surface fish and cavefish morphotypes (Figure 5). Interestingly, the testes transcriptome was more similar between morphotypes than the ovary transcriptome. We found 1,505 DEG comparing juvenile testes and 4,038 DEG comparing adult testes of Pachón cavefish versus surface fish (Figure 4A and 4B, respectively). Many of the most highly expressed DEG in either morph are unannotated, novel genes. However, the most statistically significantly upregulated gene in adult Pachón testes was Syntaxin 5a-like (*stx5al*), which is expressed in mammalian Sertoli cells (somatic nurse cells of the testicular germ cells), but has also been reported as a primordial germ cell marker in zebrafish single-cell RNAseq atlas Daniocell (Sur 2023). Gene set enrichment analysis of the genes upregulated in Pachón testes revealed Biological Process GO term categories related to angiogenesis in juveniles, and cell-cell signaling in adult testes (Figure 4C-D), likely representing morphogenesis/patterning and sperm surface interactants, respectively. Consistent with these gene expression differences, we found that in surface fish testes, blood vessels or vascular bundles were scant, while in Pachón testes most testicular tubules were associated peripherally with vascular tissue (Figure 4D). Our results at the transcriptional and morphological levels, suggest increased vascularization of the cavefish testes which could have an impact on the regulation of spermatogenesis.

**Figure 4.**
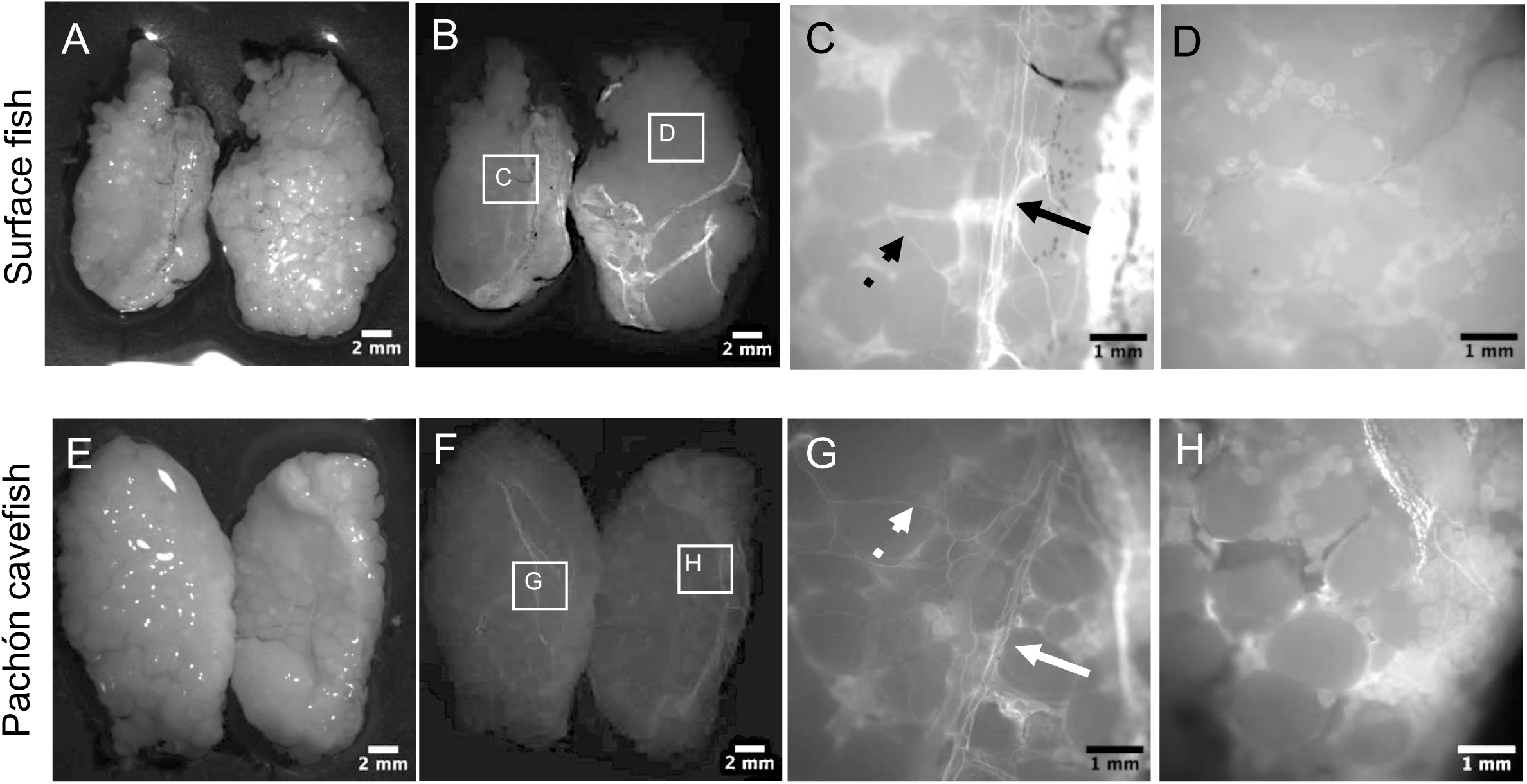
Innervation in the *Astyanax mexicanus* ovarian epithelium revealed by wholemount immunostaining. Brightfield images of the ovaries of adult surface fish (**A**) and Pachón cavefish (**B**). Lobes on the lefthand side of the image are placed with the medial side upward, while lobes on the righthand side are placed with the lateral side upward. Visualization of acetylated tubulin in wholemount fluorescence images of the ovaries shown in A, B at low magnification (**B, F**) and higher magnification (**C, D, G, H**) highlighting the differences in innervation in the medial side (**C, G**) versus lateral side (**D, H**) of the ovary. A large neuron trunk (solid arrows) can be seen on the medial side of the ovary with branches (dashed arrows) that reach throughout the ovarian epithelium.

**Figure 5.**
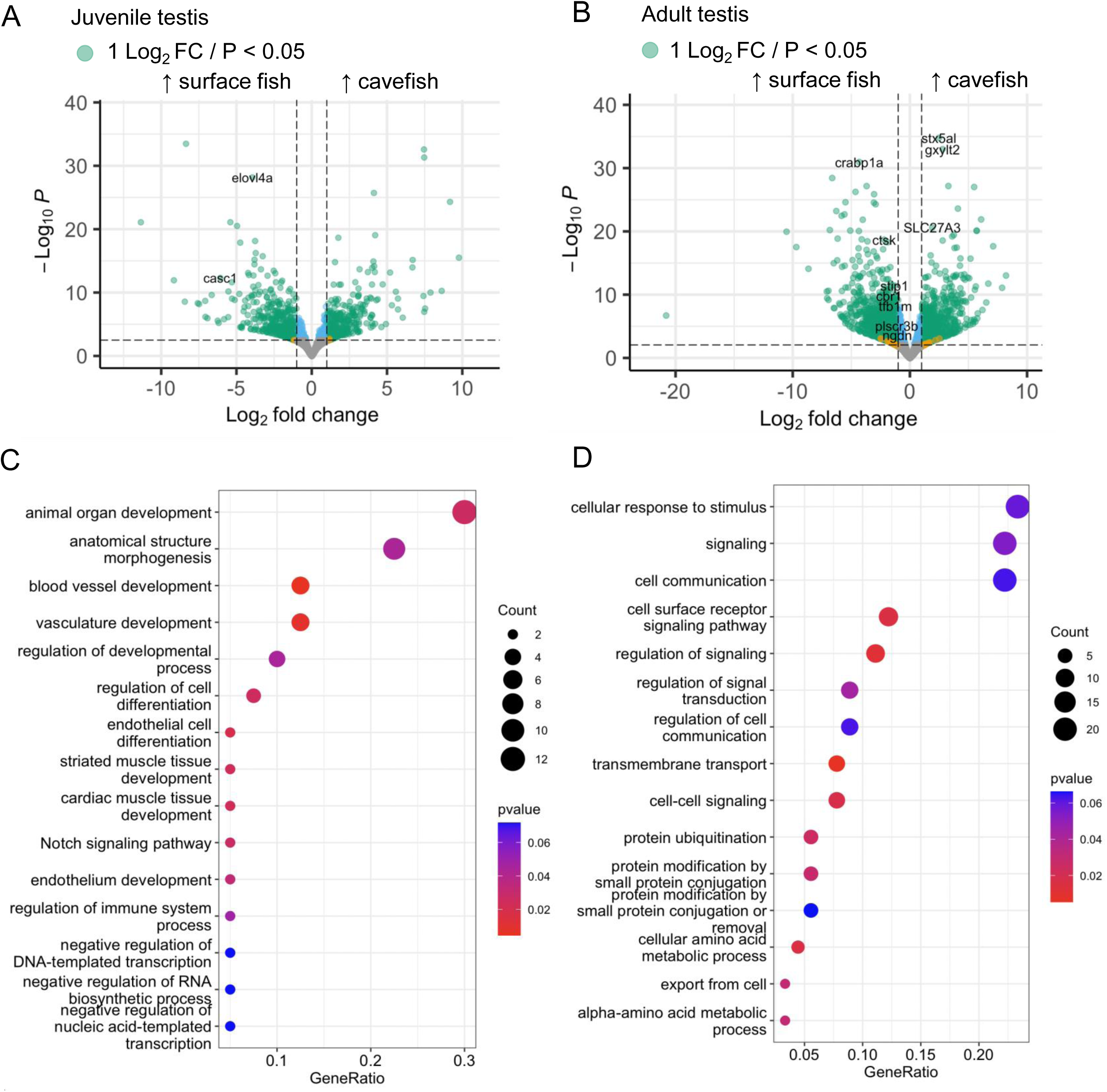
Testis transcriptome differences between surface fish and Pachón cavefish suggests enhanced angiogenesis in cavefish. (**A,B**) Volcano plots showing genes that are differentially expressed between the testes of *A. mexicanus* surface fish and Pachón cavefish at juvenile (**A**) and adult (**B**) stages. The y-axis shows −log10 adjusted p-value and the x-axis shows log2 fold change for all genes after low-count filtering. Genes with adjusted p-value < 0.05 and absolute log2 fold change > 1 are highlighted in green. Negative fold change indicates greater expression in surface fish, while positive fold change indicates greater expression in cavefish. (**C**, **D**) Dot plots displaying top 15 enriched biological process gene ontology (GO) terms identified from gene set enrichment analysis of genes upregulated in Pachón cavefish compared to surface fish at juvenile (**C**) and adult (**D**) stages. Each dot represents a biological process GO category, with the size of the dot corresponding to the number of genes in the category, and the color scale indicating the adjusted p-value. The x-axis represents the gene ratio (proportion of input genes involved in the pathway), while the y-axis lists the enriched pathways.

## Discussion

In this study, we present the first transcriptomic analysis of ovaries and testes in *Astyanax mexicanus,* comparing gene expression between lab-raised surface fish and Pachón cavefish at both juvenile and adult stages. Our results reveal clustering of samples based on developmental stage, sex, and morphotype identity, demonstrating that these factors play significant roles in shaping gonadal gene expression. This clustering supports the idea that despite the shared ancestry of surface and cave morphotypes, their reproductive systems exhibit distinct developmental trajectories resulting from evolution under different environmental pressures. This transcriptome will enable the identification and characterization of novel and conserved sex-specific genes that drive gonadal sex differentiation in juveniles and maintain sex fate determinations in adults, which in teleost fish requires both male pathway promotion and sustained female pathway suppression (Nishimura and Tanaka 2014). These findings also highlight the potential for *A. mexicanus* to serve as a model for studying the rapid evolution of highly conserved vertebrate sex regulating genes, especially in the context of extreme environmental and nutritional deficits.

### Conservation and deviation in expression profiles of conserved sex genes

Broadly compared to other male vertebrates, our *A. mexicanus* gonadal transcriptome establishes a high degree of conserved gene expression associated with testicular differentiation and spermatogenesis. All male gonad-promoting candidates investigated were expressed uniquely in, or significantly upregulated in testes of both morphotypes, including transcription factors (*dmrt1, sox9a*), signaling molecules (*amh*), and steroidogenic enzymes (*cyp11c1*). These results are consistent with expression profiles reported in other fish testis transcriptomes, and furthermore indicate homology of testicular cell subtypes relative to other vertebrates, including humans (Nynca 2022, Qian 2022, Liu 2023). However, despite typical expression of testis-enriched transcripts, our data show a striking deviation from other fishes or vertebrates with respect to the expression of canonically ovary-promoting genes. This suggests possible evolution, differential selection, or neofunctionalization of ovarian genes in *A. mexicanus*.

Most strikingly, we identified a reversal of sexually dimorphic expression in *foxl2a*, an evolutionarily conserved transcription factor produced by somatic cells of the differentiating ovary in all metazoans. Across species, *foxl2a/*Foxl2a is typically an ovary-enriched transcript and protein. Transcript levels of *foxl2a* will not necessarily correlate to protein products active in the testis, however, highlighting the importance of further experiments. Foxl2 has a complicated history of repeated gains, losses, and role changing, but generally is present in fish genomes as *foxl2a* or *foxl2b*, with some basal teleosts retaining both *foxl2a* and *foxl2b* (Bertho 2016). We identified *foxl2a*, *foxl2b*, and the paralog *foxl3* in the *A. mexicanus* ovary and testis transcriptomes. Surprisingly, our data show that *foxl2b* is more highly expressed in ovaries than testes, despite its sequence being highly divergent from other teleosts exhibiting male-enriched or non-dimorphic expression of this gene. While *A. mexicanus foxl2a* shows 75-80% identity to its teleost orthologs, there are no known studies reporting higher testicular expression of *foxl2a* in any other fishes, with most reporting low to no expression in testes (summarized in Bertho 2016). High expression of *FOXL2* protein has, however, been reported in American alligator testes prior to temperature-dependent sex determination (Yatsu 2016), which may point to latent sexual plasticity of the *A. mexicanus* testis. Taken together, it is possible that neofunctionalization – or impairment of functionalization – of *foxl2a* in *A. mexicanus* has conferred a novel role upon this gene in the testis, while *foxl2b* has retained an ovary-promoting function.

Downstream of ovary-promoting transcription factors such as *foxl2a*, expression of the steroidogenic enzyme aromatase (*cyp19a1a*) typically signals ovarian differentiation and commitment in vertebrates via production of feminizing estradiol (E2). We found both morphotypes to lack sexually dimorphic expression of this gene, suggesting deviation from typical sex hormone signaling in *A. mexicanus* and possible elevation of E2 in males. *cyp19a1a* is detected in somatic gonadal cells of both sexes, though in fish is broadly ovary-enriched and in zebrafish is required to antagonize testis fate (Devlin and Nagahama 2002, Dranow 2016). *cyp19a1a* can synthesize E2 through two biochemical pathways, a direct pathway that produces E2 using testosterone produced by the steroidogenic enzyme *hsd17b3*, and an indirect pathway that produces an estrone (E1) intermediate from androstenedione, requiring the action of *hsd17b1* to ultimately generate E2 (Miller and Auchus 2011). Our analysis revealed sexually dimorphic expression of both *hsd17b1* and *hsd17b3* (upregulated in ovaries and testes, respectively), indicating that the deployment of either E2 synthesis pathway 1) differs between sexes and 2) may ultimately result in expected rates or physiological concentrations of sex hormones. The *hsd17b1* E2 synthesis pathway is also favored in zebrafish ovaries, though has not been explored in testes, in which *cyp19a1a* expression is very low and dispensable (Liu 2022, Dranow 2016). Future studies of circulating serum hormone levels will be necessary to demonstrate whether *A. mexicanus* males exhibit physiologic aberrantly high estrogen signaling. It is also possible that male *A. mexicanus* produce an inactive *cyp19a1a* protein due to alternative splicing, as reported in the protandrous hermaphrodite barramundi (Domingos 2018).

### Upregulation of innervation in cavefish ovaries

Enrichment analysis of the DEGs upregulated in the ovaries of Pachón cavefish versus surface fish suggest possible differences in ovarian physiology. Most notably, we identified enrichment of genes involved in neuron and axon differentiation and development. This is consistent with our observation that the external ovarian epithelium is extensively innervated in both morphotypes (Figure 4). In zebrafish, the ovarian epithelium demarcates a germinal zone that is associated with nests of presumptive germline stem cells (Liu 2022). In the mammalian ovary, nerve cells are connected to the follicles as well as gonadal blood vessels (Malamed 2012), and neurotrophic growth factors and receptors are expressed at varying levels dependent on oocyte stage (reviewed in Linher-Melville and Li 2013). Given this evidence that innervation and neurotrophins influence folliculogenesis and oocyte maturation, enhanced neurogenesis in Pachón ovaries may represent an adaptive reproductive strategy that regulates renewal versus differentiation of ovarian germ cells in the cave environment, which is reproductively challenging both in terms of nutrient deprivation and mate selection.

Additionally, increased innervation may facilitate reproductive success in caves in the absence of visual stimuli, by integrating other sensory inputs from the environment that serve as cues for spawning (e.g. temperature, water currents, precipitation). In zebrafish, production of norepinephrine within neurons of the ovary is important for regulating oocyte maturation in response to nutrient cues (Kim 2021). Increased sensitivity to such cues may provide cavefish females an energy-saving advantage as nutrient availability fluctuates seasonally (Espinasa 2017). The functional significance of ovarian innervation in cavefish remains speculative, but further studies comparing oocyte development between morphotypes may elucidate how cave environments influence reproductive physiology.

### Upregulation of vascularization in cavefish testes

Enrichment analysis uncovered a striking upregulation in the expression of angiogenesis-related genes in Pachón cavefish compared to surface fish. This molecular evidence is supported by our histological findings of increased vasculature in Pachón cavefish testes (Figure 6). Vertebrate seminiferous tubules are typically avascular and form a blood-testis barrier that occludes the germinal epithelium (spermatogonia, spermatocytes, sperm, and Sertoli support cells), though specialized non-fenestrated capillaries form in the interstitial space outside of the seminiferous tubules where the steroidogenic Leydig cells reside (Ergün 2016). In mammals, upregulation of vascularization is necessary for initiating organogenesis of the testis, in particular ensuring development of fetal testis cords and limiting differentiation of the Leydig cell population (reviewed in Gu 2021). Thus, cavefish testicular vascularization may indicate a delayed or repressed state, with respect to sexual maturation. Across species, testes vascularization allows for thermoregulation, establishing optimal timing and temperature for spermiogenesis (Setchell 1998). In many fishes, temperature fluctuation in either direction lowers testosterone production and inhibits germ cell differentiation (Manning and Kime 1985, Konkal and Ganesh 2021). Testicular thermoregulation may be particularly challenging in chronically low-temperature cave environments, and the temperature preferences exhibited by cavefish could represent a reproductive strategy that preserves male fertility (Tabin 2018).

**Figure 6.**
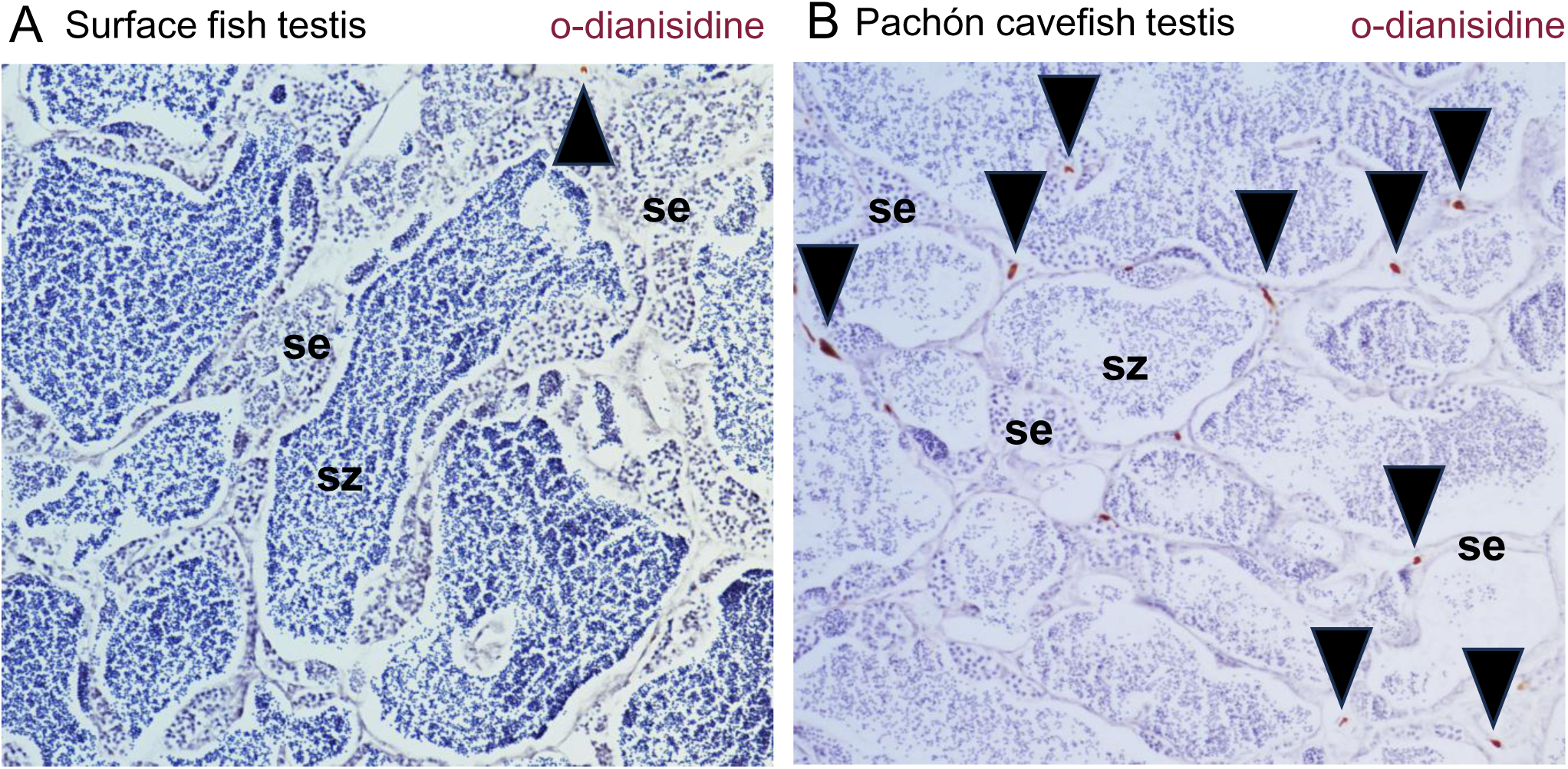
Histological comparison of testis vasculature between *Astyanax mexicanus* surface fish and cavefish. Histological sections of adult testes stained to visualize blood vessels in surface fish (A) and Pachón cavefish (B). Vasculature is stained using o-dianisidine (brown) with hematoxylin (blue) as a counterstain. Surface fish testis section (B) shows many seminiferous tubules and only a single vascular bundle (black arrow). In contrast, cavefish testis section shows numerous vascular bundles around the seminiferous tubules (black arrows). se = seminiferous epithelium; sz = spermatozoa.

Cave environments are also hypoxic, a condition known to induce angiogenesis and vascular branching via upregulated Notch signaling (Krock 2011). Pachón cavefish have adapted to hypoxia by developing more erythrocytes and enlarged hematopoietic domains, driven by an increase in Sonic hedgehog (*shh*) (van der Weele 2024). These studies align with our findings, which show upregulation of *notch1b*, *notch3*, and *shh* specifically in the Pachón testes (Additional file 2). Further comparative analysis of surface and cavefish testes will be needed to characterize differences in rate of spermatogenesis or time to sexual maturity. Our study provides molecular evidence for gonadal adaptation that allows for morphogenesis and spermiogenesis to proceed in the extremes of cave environments.

## Conclusions

Our study provides new insights into the molecular, functional, and structural differences between surface fish and cavefish gonads, gametes, and fertility, contributing to the broader understanding of reproductive biology in *A. mexicanus*. Reversal or loss of sex-specific gene expression (*foxl2a, cyp19a1a*) suggests novel functional changes or plasticity emerging in conserved sex regulating pathways in this species. The upregulation of neuron development in ovaries and increased angiogenesis in testes highlight the potential for adaptive changes in gonadal tissue architecture and function in response to environmental pressures in the cave.

These findings open several avenues for future research, including functional studies to determine how these molecular changes impact reproductive success and how environmental factors shape the evolution of sexual dimorphism and reproduction in cave-adapted populations. Overall, our results establish a foundation for further investigations into the evolution of sex-regulatory pathways and sexual dimorphism in organisms in extreme habitats.

## Declarations

### Ethics approval and consent to participate

This research was performed in compliance with ethical guidelines mandated by Harvard Medical School IACUC (Institutional Animal Care and Use Committee). All fish husbandry, euthanasia, and experimental procedures were conducted according to protocols approved by IACUC and veterinary staff (IACUC Protocol IS00001612-6 [previously identified as #04174], valid through 2026). Fish were humanely euthanized by immersion in a lethal dose (400 ppm) of tricaine (MS-222) dissolved in their environmental water, according to the approved IACUC protocol.

### Consent for publication

Not applicable

### Availability of data and materials

The RNA sequencing datasets generated and analyzed during the current study are available in the NCBI SRA repository (Accession: PRJNA1202205 [Reviewer link: https://dataview.ncbi.nlm.nih.gov/object/PRJNA1202205?reviewer=cecqungnrd9j6h5njki7u7nshh])

### Authors’ contributions

KW conceived the study, collected samples, analyzed data, and wrote and edited manuscript draft. BP collected samples, analyzed data, and edited the manuscript draft. HVG, JP, and JH processed and analyzed RNAseq data, and edited the manuscript draft. MR conceived the study, analyzed data, and wrote and edited manuscript draft.

## Acknowledgements

Brian Martineau, Emma Ferrante, and Taylor Rebbe provided fish husbandry in the aquatic facility of Clifford J. Tabin at Harvard Medical School.

## Additional material

**Additional file 1.pptx** Boxplots displaying weight of fish (A-B) and gonad (C-D) for the samples used for RNAseq analysis. Juvenile gonads were too small to accurately measure weight. In boxplots central line represents median, upper and lower edges indicate first and third quartiles, and whiskers extend to 1.5 times the interquartile range. No significant differences were observed between sample types (t-test p > 0.05).

**Additional file 2.pptx** Plots showing *A. mexicanus* gonad log2 transcript abundance of genes potentially involved in steroidogenesis (*hsd17b1* and *hsdb17b3*) or vascularization and angiogenesis (*notch1b, notch3, shh*). Each point represents the transcript abundance in an individual sample, colored and organized on the x-axis by the sample type. From left to right: cavefish (CF) juvenile ovary, CF adult ovary, surface fish (SF) juvenile ovary, SF adult ovary.

