## Supplementary figures and images for "Differential expression of sex regulatory genes in gonads of *Astyanax mexicanus* surface fish and cavefish"

### Additional file 1

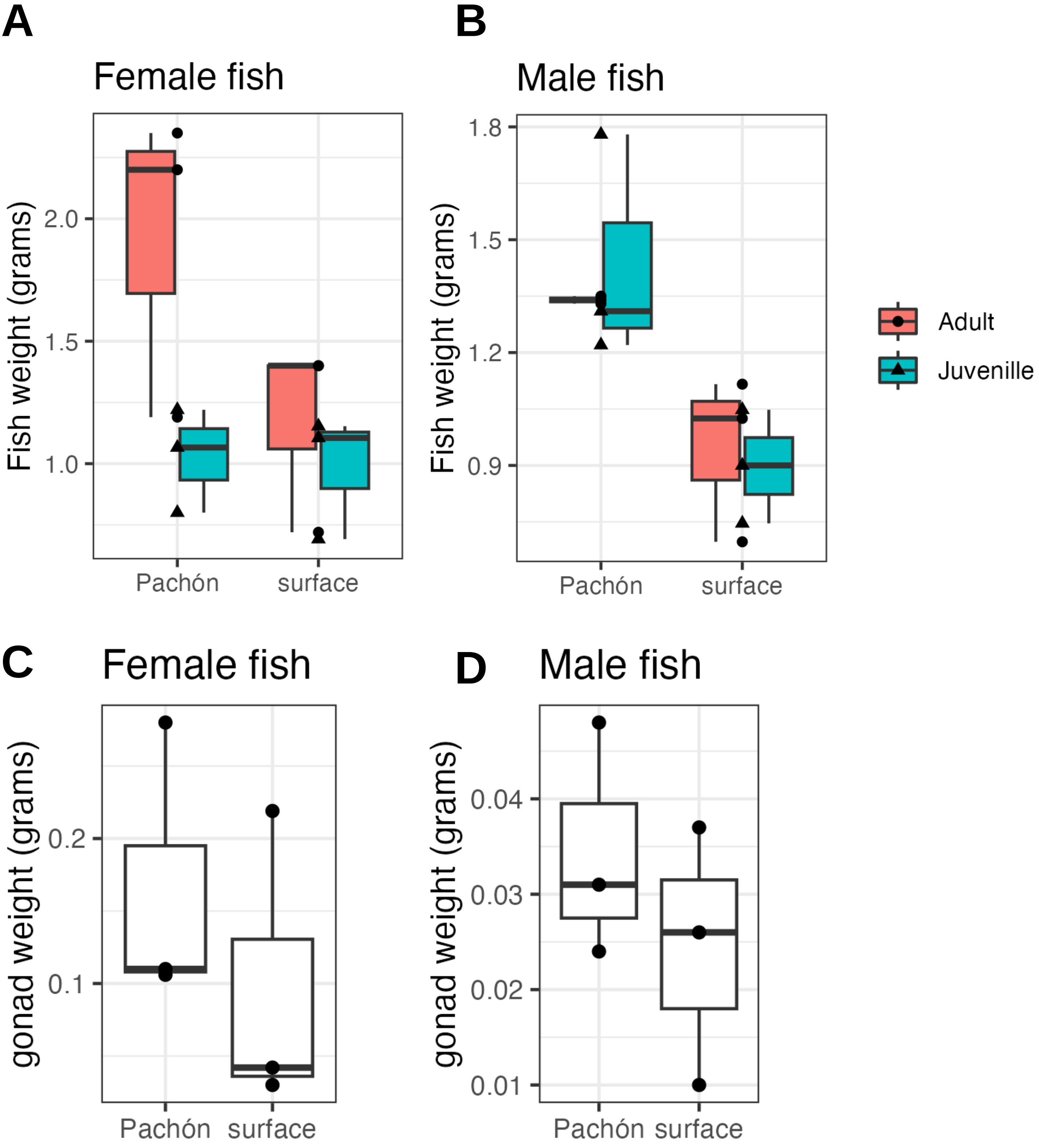

### Additional file 2

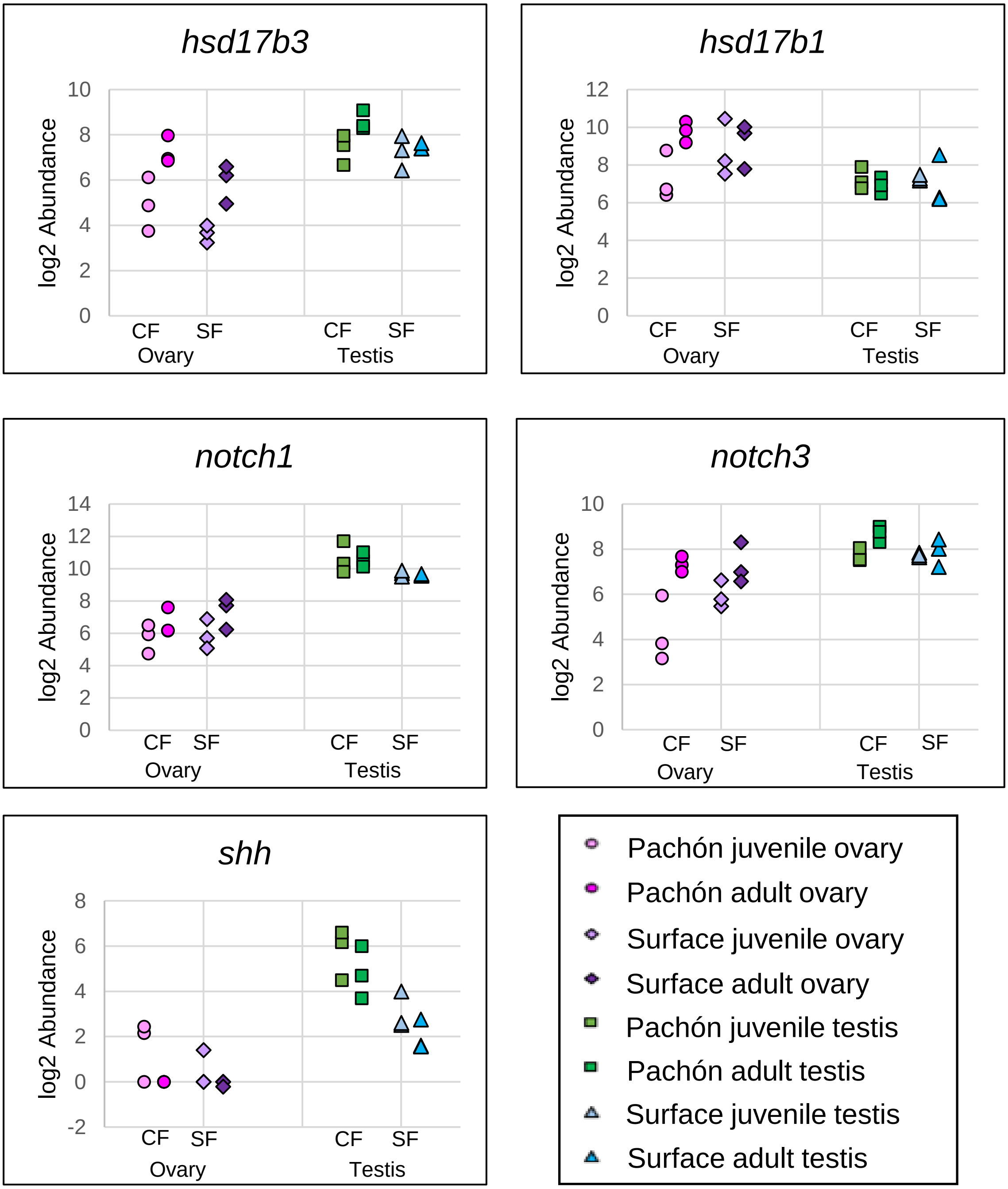
